# *Cosi2* : An efficient simulator of exact and approximate coalescent with selection

**DOI:** 10.1101/005090

**Authors:** Ilya Shlyakhter, Pardis C. Sabeti, Stephen F. Schaffner

## Abstract

**Motivation:** Efficient simulation of population genetic samples under a given demographic model is a prerequisite for many analyses. Coalescent theory provides an efficient framework for such simulations, but simulating longer regions and higher recombination rates remains challenging. Simulators based on a Markovian approximation to the coalescent scale well, but do not support simulation of selection. Gene conversion is not supported by any published coalescent simulators that support selection.

**Results:** We describe *cosi2*, an efficient simulator that supports both exact and approximate coalescent simulation with positive selection. *cosi2* improves on the speed of existing exact simulators, and permits further speedup in approximate mode while retaining support for selection. *cosi2* supports a wide range of demographic scenarios including recombination hot spots, gene conversion, population size changes, population structure and migration.

*cosi2* implements coalescent machinery efficiently by tracking only a small subset of the Ancestral Recombination Graph, sampling only relevant recombination events, and using augmented skip lists to represent tracked genetic segments. To preserve support for selection in approximate mode, the Markov approximation is implemented not by moving along the chromosome but by performing a standard backwards-in-time coalescent simulation while restricting coalescence to node pairs with overlapping or near-overlapping genetic material. We describe the algorithms used by *cosi2* and present comparisons with existing selection simulators.

**Availability:** A free C++ implementation of *cosi2* is available at http://broadinstitute.org/∼ilya/cosi2.

## 1 Introduction

A wide variety of population genetic analyses depend on efficient simulation of samples under a demographic model. The most efficient simulation method works backwards in time from the sample, first simulating the sample’s genealogy according to coalescent theory and then placing mutations on that genealogy. Because of crossing over and gene conversion, the overall genealogy for a simulated region is not a tree but a directed acyclic graph whose size grows quickly with region length and with recombination rate. To address this simulation bottleneck, a Markovian approximation was proposed and implemented ([Yang *et al*. (2014) Yang, Deng, and Niu]). However, the Markovian approximation does not readily allow for the modeling of positive selection. Simulators that do support positive selection (e.g. ([Teshima and Innan (2009) Teshima and Innan] and [Ewing and Hermisson (2010) Ewing and Hermisson])) suffer from the performance limitations of the traditional coalescent, and do not support all commonly needed features (variable genetic maps, gene conversion, structured populations) within a single framework.

Here we describe the simulator *cosi2*, which implements the standard coalescent simulation algorithm but which does so using several optimizations that greatly reduce memory and CPU requirements. The use of the original coalescent allows *cosi2* to support simulation of positive selection using the standard structured coalescent approach. For additional efficiency, the Markovian approximation can be enabled while retaining support for simulating selection. *cosi2* supports population structure, population size changes, bottlenecks and migrations. Recombination rate can be varied along the simulated region, and the program includes a utility to generate genetic maps with recombination hotspots. *cosi2* also supports a simple model of gene conversion.

**Table 1:** Simulation time comparison

| Model parameters |  |  |  | Mean simulation time per replica (s) |  |  |  |
| --- | --- | --- | --- | --- | --- | --- | --- |
| $L$ | $N$ | $k$ | $r$ | *msms* | *mbs* | *cosi2* | *cosi2* (apx) |
| 3 | 30k | 40 | 1e-8 | 3 | 76 | .7 | .25 |
| 10 | 30k | 80 | 1e-8 | 40 | > 1000 | 5 | 1 |
| 30 | 30k | 200 | 2e-8 | 2786 | > 10000 | 162 | 21 |
| 60 | 25k | 200 | 2e-8 | 10286 | > 20000 | 686 | 71 |
| 3 | 30k | 40 | 1e-8;hs | n/s | 69 | .76 | .26 |
| 10 | 30k | 80 | 1e-8;gc | n/s | n/s | 9 | 3 |

Model parameters: $L$ - region length in megabases, $N$ - diploid effective population size, $k$ - sample size, $r$ - recombination rate (hs - with hotspots, gc - with gene conversion). Common parameters: mutation rate $m = 1e - 8$ , selection coefficient $s = .0185$ , present-day frequency of selected allele $f = .7$ . n/s means not supported. *cosi2* (apx) is *cosi2* with Markov approximation.

## 2 Algorithm description

Here we describe the different optimizations which, taken together, greatly improve the scaling behavior of coalescent simulations. We also describe the implementation of positive selection modeling in *cosi2*. Additional details of the algorithms are given in the Supplemental Information.

### 2.1 Tracking only the veneer of the Ancestral Recombination Graph

Previously described coalescent simulators work in two phases. First, they simulate the Ancestral Recombination Graph (ARG) representing the coalescence and recombination events in the history of the sample. Second, they place mutations on edges of the ARG, tracing down the graph to find the leaves inheriting each mutation. *cosi2* combines these into a single phase: mutations are placed on an ARG edge as soon as the edge is created by a coalescence or a recombination; the edge and the child node at its end are then immediately discarded. Thus, only a thin veneer of the ARG is kept at each point of the simulation, consisting of ARG nodes which do not yet have a parent. This greatly reduces memory requirements and improves memory access locality, but creates a problem when placing mutations: determining the set of leaves inheriting a mutation can no longer be done by tracing down the ARG. *cosi2* solves this by keeping, for each chromosome segment carried by an ARG node, the set of leaves (the leafset) inheriting that segment. Leafsets are updated incrementally at each simulation step.

### 2.2 Efficiently representing segment lists

When representing an ARG, a node needs to record only those segments of each chromosome that will have a descendent in one of the final, sampled chromosomes. With extensive recombination, the segments become highly fragmented and efficient handling of the segment list is crucial to good performance. In *cosi2*, since nodes participating in recombination or coalescence are discarded immediately after the operation, their segment lists can be used as building blocks for segment lists of the recombined or coalesced nodes. Representing the segment lists with a modified skip-list randomized data structure ([Pugh (1990) Pugh]) permits all segment list operations to run in logarithmic time. By keeping additional information in the skip-list, finding the location of the next mutation during mutation placement also becomes efficient.

### 2.3 Efficient sampling of crossovers and gene conversions

The vast majority of crossover and gene conversion events do not change the ARG but do take time to track. These are either crossovers falling entirely to one side of a node’s segment list, or gene conversions falling entirely outside any segment. *cosi2* directly samples only crossovers and gene conversions that actually change the ARG, by keeping the crossover and gene conversion rates of the individual nodes in a data structure supporting quick access to the total rate and efficient sampling of event location weighted by the genetic map. The skip-list representation of segment lists allows incremental updating of the node’s individual event rates.

### 2.4 Modeling of selection

Selection is implemented using the structured coalescent approach ([Teshima and Innan (2009) Teshima and Innan]). At start of simulation, sampled chromosomes are partitioned into two classes based on their allelic state at the selection site. Coalescence happens only within the same allelic class, with coalescence rate based on the frequency trajectory of the causal allele. The frequency trajectory of the selected allele can be deterministic (based on causal allele age and present-day frequency), or can be provided externally; the latter mode permits the use of stochastic trajectories.

### 2.5 Implementing the Markovian approximation

We implement an approximation to the coalescent, in which coalescences are restricted to occur between nodes whose genetic information overlaps or nearly overlaps ([McVean and Cardin (2005) McVean and Cardin]). By modifying only the coalescence step within an existing coalescent simulator that supports selection, we add the ability to approximate the coalescent while preserving support for simulating selection. The key algorithmic difficulty is in determining the number of coalesceable node pairs, and in choosing a pair uniformly at random from all such pairs. We maintain this information incrementally using a dynamic data structure based on augmented interval trees.

For each ARG node *n*, let [*n^b^, n^e^*] be the range of the simulated region covered by the node’s genetic material. We define a *hull* of *n* as the region [*n^b^, n^e^* + *u*] where *u* is a parameter controlling the amount of approximation. Two nodes are coalesceable if their hulls overlap; *u* therefore specifies the maximum separation between coalescing nodes. When a hull [*n^b^, n^e^*] is added, we can quickly find the number of hulls it overlaps by subtracting from the total number of hulls the count of those ending before *n^b^* or starting after *n^e^* ([Layer *et al*. (2013) Layer, Skadron, Robins, Hall, and Quinlan]). Hull removal is analogous. Since all coalescent operations (crossover, gene conversion, coalescence and migration) can be implemented as a sequence of hull additions and removals, we can maintain the count of coalesceable node pairs with only logarithmic overhead. Selecting a coalesceable pair uniformly at random requires maintenance of additional information; description of the implementation – also requiring only logarithmic time and space overhead – is given in Supplemental Information.

## 3 Summary

*cosi2* provides a combination of performance and supported demographic scenarios unavailable with existing selection-supporting simulators. We hope it becomes a useful tool for population geneticists studying positive selection.

## Acknowledgement

**Funding:** PCS is funded by an NIH Innovator Award 1DP2OD006514-01 and by a Broad Institute SPARC award.

